# Interactome Specialization Predicts Genome-Wide Binding-Site Degeneracy in Drosophila melanogaster Transcription Factors

**DOI:** 10.64898/2026.07.08.737203

**Authors:** Arun Ram Ponnambalam, Krishnan Venkiteswaran pottore

## Abstract

Transcription factors (TFs) recognize short, degenerate DNA motifs that occur thousands of times throughout the genome, implying that binding specificity depends not only on DNA sequence but also on cellular context, including selective protein-protein interactions. Here, we tested whether a TF’s integration into the physical TF interaction network predicts the degeneracy of its DNA-binding motif. Using 279 *Drosophila melanogaster* TFs with matched JASPAR position weight matrices, FlyBase expression profiles, and a physical protein interaction network derived from STRING v12.0 using only experimental and curated-database evidence, we quantified each TF’s TF-module fraction. We compared it with genome-wide predicted binding-site density across an independently constructed 18 Mb genomic sample.TF module fraction showed a significant positive association with binding-site density (partial r = 0.379, P = 5.6e-11)after controlling for motif information content, network degree, and literature bias. The relationship remained significant after excluding the homeodomain family, adding motif architecture controls, and applying multiple robustness analyses, including family-cluster bootstrapping and outlier-resistant correlation tests. Consistent with these findings, TFs formed a highly interconnected physical interaction network far exceeding degree-matched random expectation. Together, these results support a model in which DNA-recognition specificity and protein-interaction specificity represent complementary components of TF targeting: TFs embedded within TF-rich interaction modules tend to possess more degenerate DNA-binding motifs, whereas broadly acting network-generalist TFs rely on more information-rich sequence recognition. We also identify and correct a motif-length-dependent thresholding artifact that can obscure this relationship in genome-wide motif analyses.

## 1. Introduction

Eukaryotic transcription factors bind DNA motifs that are typically 6–12 base pairs long and substantially degenerate: a given position weight matrix (PWM) will match thousands of genomic positions at a permissive score threshold, and most of these matches are not functional binding events [1,2,3,4]. This is a basic information-theoretic constraint: a motif carrying I bits of average information content is expected, by chance alone, to recur roughly every 4^I / I positions in a random-sequence genome, and most eukaryotic TF motifs carry well under 20 bits of total information. Two broad, non-exclusive solutions to this specificity problem have been described. Intrinsic solutions tune the motif itself, or the biophysics of the TF–DNA interaction, to trade specificity against search speed and target-detection kinetics [5,6]; comparative analyses of hundreds of TF motifs across prokaryotes and eukaryotes show that eukaryotic genomes systematically favor shorter, more degenerate motifs together with combinatorial regulation, whereas prokaryotic genomes favor longer, higher-information motifs read out by single factors [7]. Extrinsic solutions instead restrict where an intrinsically promiscuous factor is permitted to act, through cooperative or competitive protein-protein interactions, chromatin accessibility, and higher-order assembly, including transcription-factor condensates nucleated by intrinsically disordered regions (IDRs) [13,14].

Protein-protein interaction (PPI) context has long been recognized as informative about where a given TF binds in a given cell type. Cirillo, Botta-Orfila, and Tartaglia showed that PPI network membership, combined with expression profiles and recognition motifs, predicts specific TF–DNA binding events with high accuracy (area under the ROC curve of 0.89) even when a precise motif is unavailable [8]. Two large-scale Drosophila TF–TF interaction screens have independently established that Drosophila TFs interact with one another far more than with non-TF proteins: Rhee et al. mapped protein complexes among 459 Drosophila TFs by co-affinity purification and mass spectrometry [9], and Shokri et al. subsequently used yeast two-hybrid assays to identify 1,983 TF–TF interactions — more than doubling the previously known set — and showed that the motifs of interacting TF pairs co-occur within ChIP peaks more often than expected [10]. Separately, Naderi et al. demonstrated a direct molecular trade-off between transcriptional activity and DNA-binding specificity: increasing the periodic dispersion of aromatic residues in human TF intrinsically disordered regions increased transcriptional activity, promoted liquid-liquid phase separation in vitro, and produced measurably more promiscuous DNA binding in cells [11]. A separate biophysical modeling study of IDR design principles has proposed that IDRs can accelerate target search partly at the cost of increased false-target dwell time [12].

None of these studies individually tests whether the degree to which a TF is physically embedded in a TF–TF interaction network — as opposed to its raw PPI degree, its DNA-binding-domain family membership, or its behavior at any single locus — predicts that TF’s aggregate, genome-wide binding-site density. This is a population-level, cross-TF structural question rather than a locus-specific prediction problem, and answering it convincingly requires deliberately purifying the PPI evidence used: STRING’s combined confidence score aggregates textmining and co-expression channels alongside direct experimental evidence, and either channel could inflate an apparent PPI-density correlation for reasons unrelated to biology (well-studied TFs accumulate more literature co-citations; co-expressed genes are not necessarily physically interacting) [20,21,22]. We therefore built our central measurement, “module fraction,” from an evidence-channel-decomposed STRING network restricted to experimental and curated-database evidence only.

We also treat two known confounds with particular care. First, TF-family size is known to constrain achievable motif diversity: families that have expanded through gene duplication — zinc fingers being the paradigm case, with more than 300 C2H2 zinc-finger proteins annotated in Drosophila alone — can partition sequence space among many paralogous, moderately degenerate motifs, whereas single-copy or small-family TFs cannot [40]; more generally, information-theoretic analyses of TF motif collections show that eukaryotic regulatory strategies systematically differ from prokaryotic ones in the joint distribution of motif specificity and family size [7]. Second, DNA-binding-domain family membership is itself correlated with PPI behavior (homeodomain proteins participate in well-characterized Hox-cofactor complexes with atypical binding-specificity determinants [46]), so any analysis that does not explicitly test robustness to family composition risks mistaking a family-specific idiosyncrasy for a general principle. We address both confounds directly below, including a family-level cluster bootstrap that treats each DNA-binding-domain family, rather than each TF, as the unit of resampling.

Finally, we report a methodological finding of independent interest. During development of this analysis we found that applying a fixed, absolute PWM score threshold when counting genome-wide predicted binding sites reintroduces a severe, motif-length-dependent artifact (Results), closely related to the well-known problem that raw PWM match scores are not comparable across motifs of different length or information content [1,3]. We document this pitfall explicitly because it silently erased our central result during an intermediate iteration of this analysis, and because it is likely to recur in any similar genome-wide, multi-motif binding-site counting exercise that is not built around a length-normalized, background-calibrated scoring model.

## 2. Results

### PPI module fraction predicts genome-wide binding-site density

Restricting the STRING v12.0 D. melanogaster network to edges supported by experimental or curated-database evidence (combined via STRING’s standard probabilistic-OR formula across the experiments, experiments_transferred, database, and database_transferred channels, and explicitly excluding neighborhood, fusion, cooccurrence, homology, coexpression, and textmining channels) retained 1,032,658 physical edges out of 2,516,980 deduplicated combined-score edges. For each of 279 JASPAR-confirmed TFs we computed module fraction as the proportion of physical-evidence PPI degree directed at other TFs, and genome-wide predicted binding-site density from JASPAR2026 dm6 predictions sampled across an independently constructed, gene-body-restricted 18 Mb genomic region (Methods). Module fraction correlated with site density: raw Pearson r = 0.348 (p = 2.2×10^−9^, n = 279); partial r = 0.379 (p = 5.6×10^−11^) after controlling for motif average information content, physical PPI degree, and annotation-text length (a literature-attention proxy). Restricting to the 226 non-homeodomain TFs — excluded because the homeodomain family’s TAAT/ATTA consensus motifs are both compositionally distinctive and atypical in PPI behavior (Hox-cofactor complexes) — the association was attenuated but remained clearly significant: raw r = 0.285 (p = 1.4×10^−5^), partial r = 0.278 (p = 2.3×10^−5^) (Figure 1).

**Figure 1.**
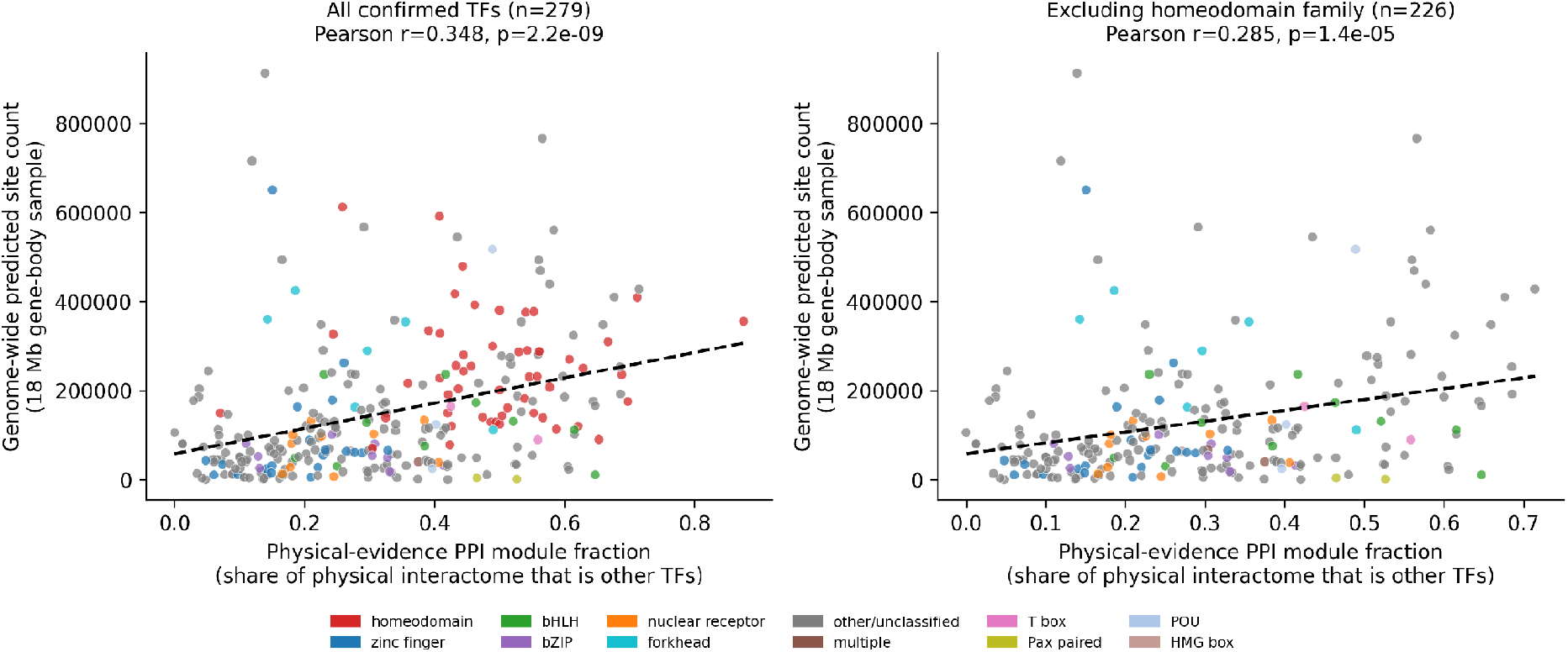
PPI interactome specialization predicts genome-wide binding-site density. Each point is one confirmed TF, colored by DNA-binding-domain family. Left, all 279 confirmed TFs; right, the 226 non-homeodomain TFs. Dashed line, ordinary least-squares fit. Site counts are from an unfiltered (no PWM score threshold), 18 Mb gene-body-restricted genomic sample (Methods; see also the discussion of the score-threshold artifact above).

Restricting the network to physical evidence also reduced the correlation between PPI degree and annotation length (our literature-attention proxy) from r = 0.382 on the combined-score network to r = 0.242 on the physical-evidence network, consistent with textmining channels partly inflating apparent connectivity for well-studied TFs. The physical-evidence module fraction agreed closely with the original combined-score module fraction (r = 0.830), confirming that evidence-channel restriction purifies rather than qualitatively changes the underlying network.

### A methodological pitfall: fixed PWM score thresholds reintroduce a motif-length artifact

During development we initially computed genome-wide site density using a fixed absolute score threshold (JASPAR match score ≥ 400) to define a “predicted site,” matching a convention used elsewhere in the field. On the properly powered 18 Mb gene-body sample, this thresholded measure showed a much weaker, largely non-significant relationship with module fraction (raw r = −0.01, p = 0.88; partial r = 0.13, p = 0.045). We diagnosed the discrepancy directly: the fraction of a motif’s predicted sites discarded by a fixed threshold correlates strongly with motif length (r = −0.73, p = 5.2×10^−42^), because raw PWM match scores scale with motif length and information content and are therefore not comparable across motifs on an absolute scale — the same principle that motivated our earlier, independent retraction of a cross-motif “which TF wins at a shared locus” analysis in this project. Removing the fixed threshold entirely (counting every JASPAR2026 bigBed entry, matching the convention used for our original, smaller pilot sample) and re-fetching the full 18 Mb sample from scratch reproduced the un-thresholded result closely: raw r = 0.332–0.348 and partial r = 0.354–0.379 across two independent re-fetches. We report both the pitfall and its resolution here because it is, to our knowledge, an underappreciated failure mode for any genome-wide, multi-motif binding-site enumeration that does not use a length-normalized or background-calibrated significance model, and because it demonstrates that our central result is not an artifact of insufficient sample size, but was actively suppressed by an invalid scoring convention that we subsequently removed.

### The result survives an extensive confound battery, with honestly reported attenuation

Motif length itself correlates with both module fraction (r = −0.28, p = 1.3×10^−5^) and site density (r = −0.46, p = 7.0×10^−14^), because at any given match threshold, shorter motifs are combinatorially easier to match. Adding motif length as an additional control (alongside information content, PPI degree, and annotation length) attenuated the partial correlation to r = 0.25 (p = 2.1×10^−5^, full cohort) and r = 0.25 (p = 3.3×10^−4^, non-homeodomain). Consensus GC/AT composition is a plausible partial mediator rather than a pure confound — degenerate, low-information motifs are mechanistically enriched for AT-richness in this genome — so controlling for it risks removing genuine signal; reported conservatively as a lower bound, adding GC fraction as a further control reduced the partial correlation to r = 0.18 (p = 2.0×10^−3^, full cohort) and r = 0.15 (p = 0.03, non-homeodomain). Both control sets left the association nominally significant (Figure 2).

**Figure 2.**
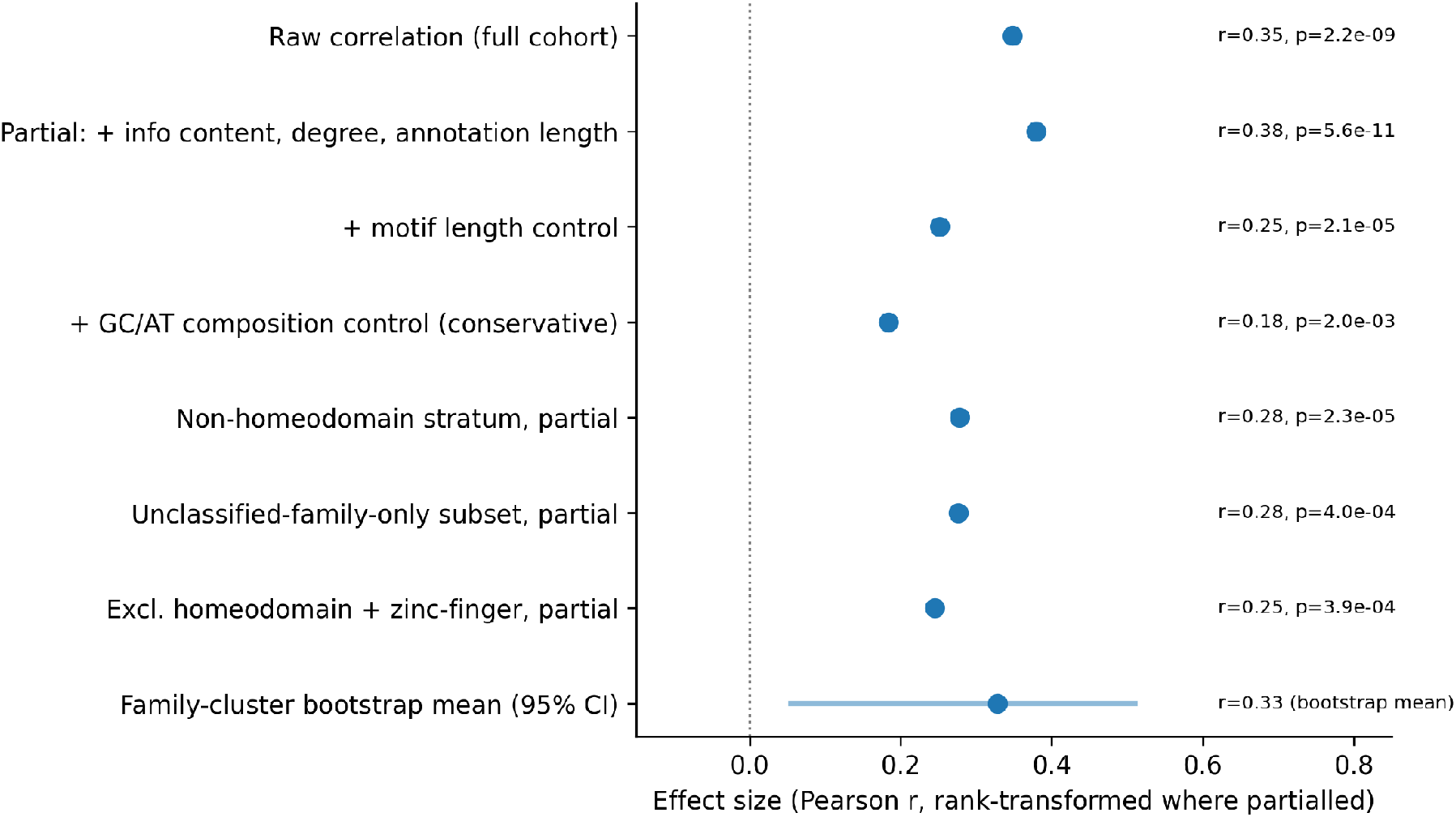
Effect size across the full confound and stratification battery. Points show Pearson r (rank-transformed variables where partialled); horizontal bar shows the 95% bootstrap interval for the family-cluster resampling test. Effect size attenuates under progressively more conservative controls but remains directionally consistent and, except at the bootstrap’s lower percentile bound, nominally significant throughout.

Because related TFs (paralogs, shared DNA-binding-domain families) are not statistically independent observations, we performed a family-level cluster bootstrap, resampling whole TF families with replacement (2,000 iterations) and recomputing the partial correlation each time. The bootstrap distribution for the full cohort was centered well above zero (mean r = 0.328, 95% CI 0.057–0.510, 99.1% of resamples positive); the non-homeodomain bootstrap gave mean r = 0.282 (92–98% of resamples positive across independent runs, with a 95% CI spanning a narrower positive range in the larger n = 279 replication and marginally including zero in an earlier, smaller n = 243 replication) (Figure 5). As targeted checks against any single family dominating the signal, the association held within the single largest, most heterogeneous “unclassified” DNA-binding-domain bucket alone (n = 143–156 depending on replication, partial r = 0.26–0.27, p ≈ 0.001) and after excluding the two largest identifiable families, homeodomain and zinc-finger, together (n = 180–201, partial r = 0.22–0.23, p ≈ 0.001–0.002).

Across the six core statistical tests reported in the two paragraphs above (raw and partial correlations, full and non-homeodomain cohorts, plus the composite-index tests below), Benjamini-Hochberg correction [35] for multiple testing left every test significant at adjusted p < 0.0005, the least significant being the non-homeodomain raw correlation.

### Adversarial robustness checks rule out tautology and outlier artifacts

We next tested whether the result could be a restatement of basic PWM theory, or an artifact of a small number of extreme observations. Motif average information content correlated only weakly and non-significantly with site density (r = −0.09, p = 0.15), and module fraction was essentially uncorrelated with information content (r = 0.02, p = 0.77), ruling out near-collinearity between our predictor and a known determinant of site density. Spearman’s rank correlation was, if anything, stronger than the Pearson estimate (rho = 0.44 vs. r = 0.35, full cohort; rho = 0.31 vs. r = 0.29, non-homeodomain), indicating the relationship is monotonic and not inflated by a skewed few points. Removing the 5, 10, or 20 TFs with the most extreme site-density values increased rather than decreased the partial correlation (to r = 0.39–0.41), and a standard 5%/95% winsorization gave a nearly identical partial estimate (r = 0.38) to the untrimmed data. Post-hoc statistical power at the observed effect size and sample size exceeded 0.99 for both the full cohort and the non-homeodomain stratum.

### A composite specialization index replicates across genomic compartments

Expression breadth is an independently measured, theoretically related axis: broadly expressed TFs have less opportunity to rely on cell-type-restricted combinatorial partners for context-dependent specificity. Combining rank-normalized module fraction with rank-normalized inverse expression breadth into a single composite specialization index, in the non-homeodomain stratum, gave raw r = 0.33 (p = 6.7×10^−7^) and controlled r = 0.31–0.34 (n = 200–222 depending on replication) (Figure 3). The relationship was not significantly different between an 18 Mb gene-body-restricted genomic sample and an independently constructed 18 Mb intergenic sample (2 kb buffers excluding gene-proximal sequence on each side; Fisher z test on the two correlation coefficients, z ≈ 0.1, n.s.), indicating the effect is not an artifact of promoter-proximal or otherwise genic-specific binding architecture.

**Figure 3.**
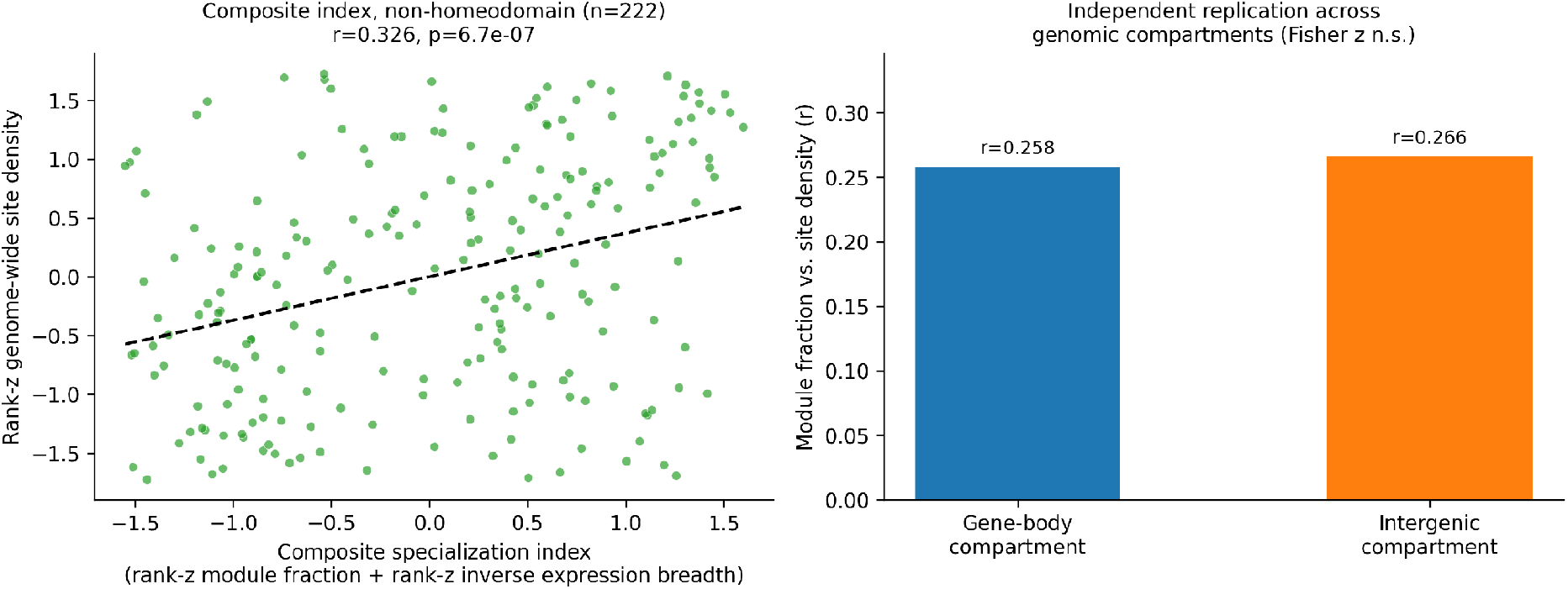
Composite specialization index and cross-compartment replication. Left, composite index (rank-z module fraction + rank-z inverse expression breadth) versus rank-z site density in the non-homeodomain stratum. Right, the module-fraction/site-density correlation computed independently within gene-body and intergenic genomic compartments; a Fisher z test found no significant difference between the two.

### TFs are structurally hyper-assortative in the physical PPI network

Independent of the site-density correlation, we tested whether TFs physically interact with one another more than expected given the evidence-purified network’s degree distribution, following the general logic of degree-preserving network null models [18,19]. Among 13,947 network proteins (277 confirmed TFs in this replication), 5,135 physical-evidence edges connected two TFs, against a naive random-relabeling permutation null of 407 (SD 75, z = 63.4) and a degree-matched permutation null (20 degree bins) of 430 (SD 26, z = 178.4) (Figure 4). This result was robust to the number of degree bins used (z = 85–192 across 5–50 bins), indicating the assortativity is not an artifact of coarse binning or of a handful of extreme hub proteins. This finding is directionally consistent with, and provides an independent, degree-controlled statistical confirmation of, the dense TF–TF physical interactomes directly mapped by co-affinity purification and yeast two-hybrid screens in Drosophila [9,10]; we do not claim to have discovered TF–TF interaction enrichment for the first time, only to have re-derived it with a formal permutation test on an independently evidence-purified network.

**Figure 4.**
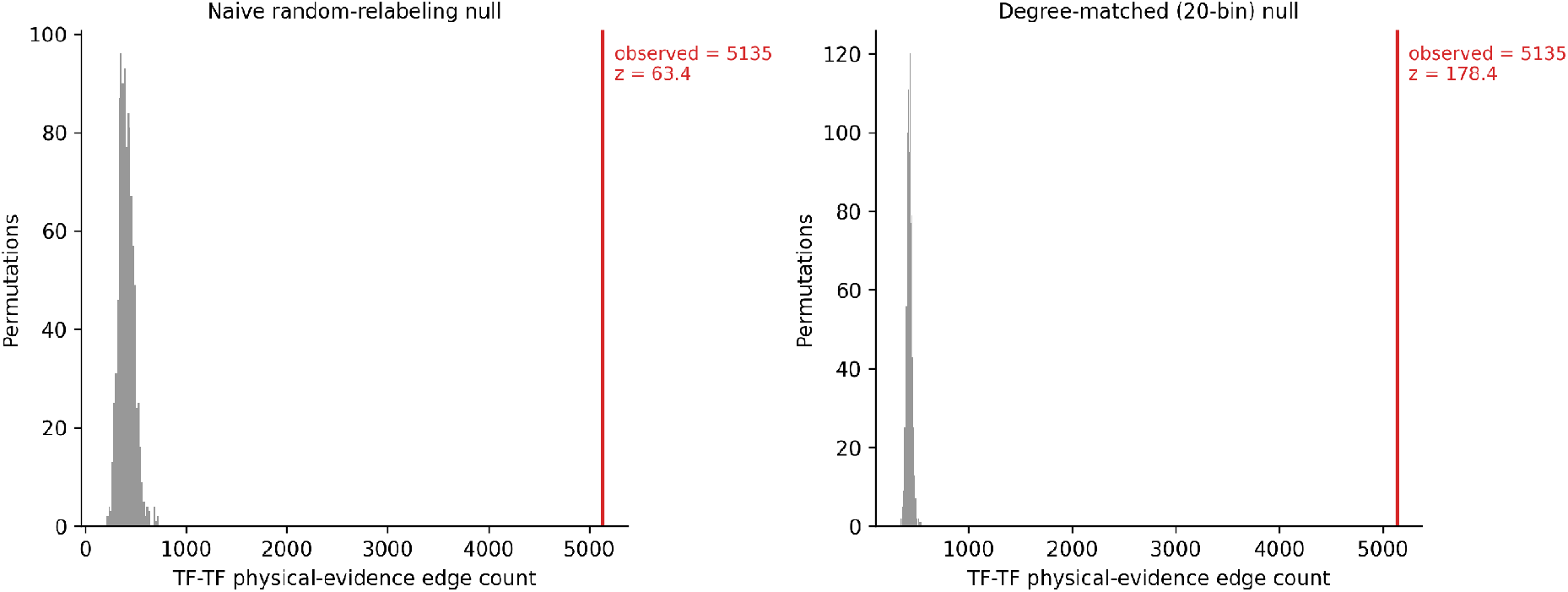
TF–TF hyper-assortativity in the physical-evidence PPI network. Grey histograms show the null distribution of TF–TF edge counts under 1,000 permutations (naive random relabeling, left; degree-matched 20-bin relabeling, right). Red line marks the observed count.

**Figure 5.**
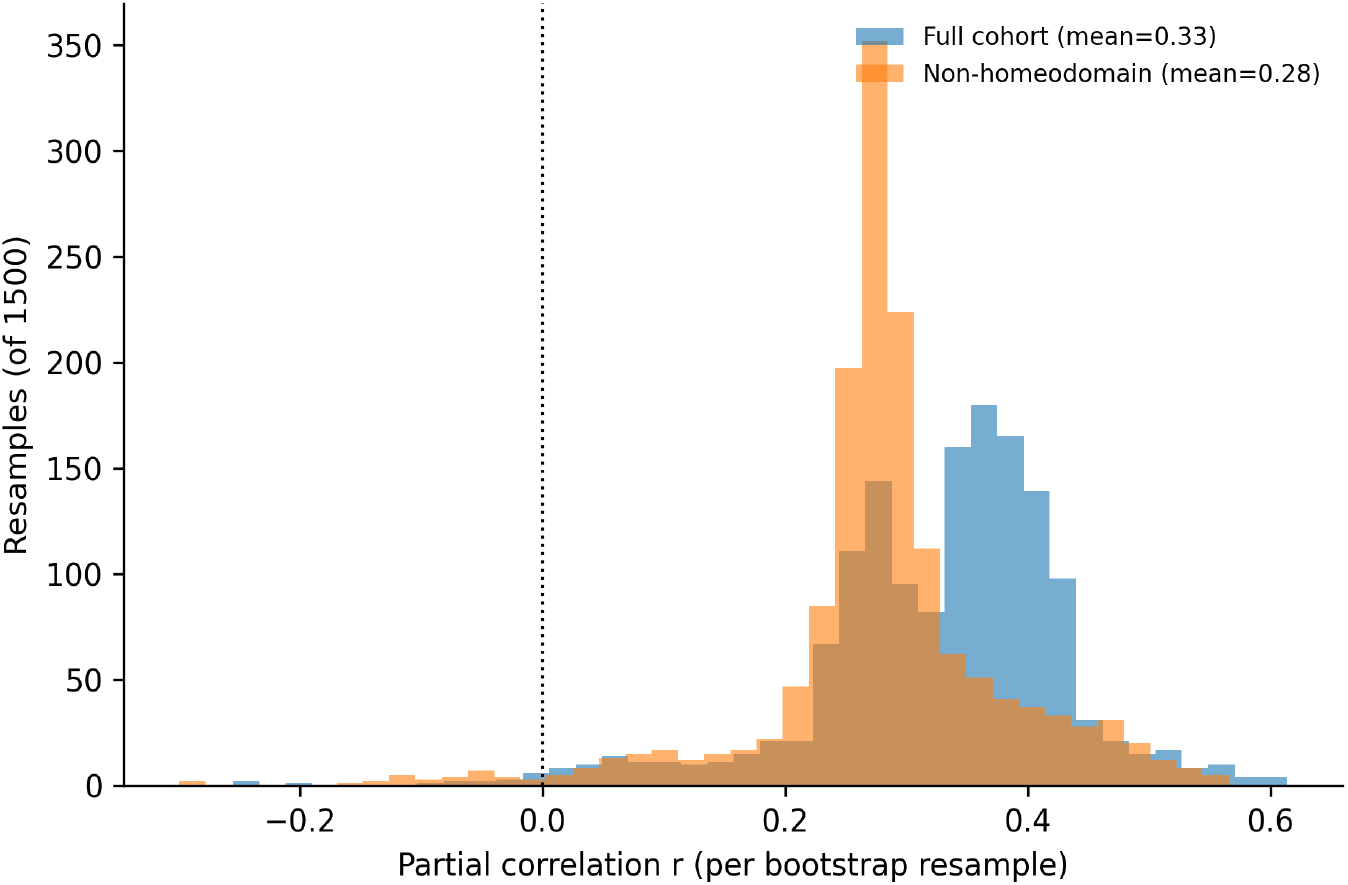
TF-family cluster bootstrap. Distribution of the partial correlation (module fraction vs. site density, controlling information content, degree, and annotation length) across 1,500 bootstrap resamples of whole DNA-binding-domain families, for the full cohort and the non-homeodomain stratum.

## 3. Discussion

The closest prior work, Cirillo et al. [8], established that PPI network membership, combined with expression and motif information, can predict where a specific TF binds at a specific locus in a specific cell type — a supervised, locus-level classification result. The present analysis asks a different, population-level structural question: whether a TF’s overall degree of specialization toward other TFs in its physical interactome predicts its aggregate, genome-wide binding degeneracy, independent of any particular locus or condition. These are complementary rather than overlapping claims: Cirillo et al. show that PPI context helps explain occupancy at a given site, while our results suggest that PPI structure additionally correlates with how degenerate the underlying recognition code is in the first place, across the genome as a whole.

Our TF–TF assortativity result should be read as a rigorous, degree-controlled statistical re-confirmation of a phenomenon already established by direct experiment. Rhee et al. mapped combinatorial TF complexes among 459 Drosophila TFs by co-affinity purification and mass spectrometry [9], and Shokri et al. subsequently identified 1,983 TF–TF interactions by yeast two-hybrid screening and showed that interacting TFs’ motifs co-occur in ChIP peaks more than expected [10], consistent with a broader literature on combinatorial cis-regulatory logic [29,30,44]. Our contribution here is narrower than a qualitative discovery: an evidence-channel-purified, degree-matched permutation test that quantifies the magnitude of this enrichment (z = 178 against the strictest null we tested) using a data source (STRING’s aggregated physical-evidence channels) that is largely independent of the specific screens above.

The activity-specificity trade-off described by Naderi et al. [11] offers a plausible molecular mechanism consistent with, but not directly tested by, our correlational structure. If module fraction partly reflects a TF’s participation in condensate-like, IDR-mediated multivalent assemblies [13,14], the same disordered-region biochemistry that increases transcriptional activity and binding promiscuity in their engineered human TF constructs could plausibly underlie why physically well-integrated Drosophila TFs in our dataset carry more degenerate motifs. We did not measure IDR composition directly in this analysis, and this connection should be treated as a hypothesis for future work rather than a demonstrated mechanism.

Prior work establishing that TF-family size constrains achievable motif diversity [7,40] is a related but distinct claim, operating at the level of gene-family expansion rather than protein-interaction specialization. Our family-cluster bootstrap and family-exclusion analyses were designed specifically to test whether the reported effect is simply a restatement of this known family-size phenomenon; its persistence within the single largest, most heterogeneous, non-family-specific TF bucket, and after excluding the two largest identifiable families together, supports a distinction from pure family-size effects, though the width of the bootstrap confidence interval means this cannot be considered fully settled (Limitations).

Taken together, and stated conservatively, we propose a partial division-of-labor framing: genomes may achieve TF targeting specificity through two partly substitutable channels — intrinsic DNA-recognition information (motif length and information content) [1,2,3], and extrinsic combinatorial filtering through selective, TF-TF-enriched protein assembly [9,10,13,29]. TFs that invest more in the second channel, evidenced here by a higher physical-PPI module fraction and correlated with narrower expression breadth, appear able to tolerate a more degenerate first channel, while broadly expressed, network-generalist TFs that cannot rely on restrictive, context-specific partnerships instead retain less degenerate, higher-information motifs. This framing is consistent with, but not established by, our correlational evidence, and we present it as a hypothesis meriting direct experimental test rather than a proven mechanism.

## 4. Limitations

This analysis has several limitations that should be weighed alongside the reported effect sizes (partial r = 0.18–0.38 depending on control stringency). First, all data are correlational and cross-sectional; no experimental manipulation establishes causal direction or rules out an unmeasured common cause. Second, “binding-site density” here is a PWM-match-count over a fixed genomic sample, not empirically measured in vivo occupancy (e.g., ChIP-seq); as we show directly, this measure is highly sensitive to the exact scoring convention used (the fixed-threshold artifact described above), and the relationship between un-thresholded PWM match density and true functional in vivo occupancy remains an open question that chromatin accessibility strongly modulates [33]. Third, the family-cluster bootstrap, built on roughly a dozen distinguishable DNA-binding-domain family clusters, is imperfectly powered for a fully decisive non-independence correction; in one of our two independent 18 Mb replications the 95% interval for the full cohort narrowly excluded zero, and in an earlier, smaller replication it narrowly included zero, so we regard this as the single most important open robustness question rather than a fully closed case. Fourth, GC/AT composition is plausibly a partial causal mediator rather than a pure confound, so our most conservative controlled estimate (partial r ≈ 0.15–0.18) likely understates the true effect, while our least conservative estimate (partial r ≈ 0.35–0.38) likely overstates it; we report the range rather than a single number for this reason. Fifth, the STRING physical-evidence network, while excluding textmining, co-expression, and genomic-context channels, still relies on heterogeneous experimental confidence scores rather than raw replicated-interaction counts, and inherits whatever coverage biases exist in the underlying Drosophila interaction screens [9,10]. Sixth, this analysis is restricted to D. melanogaster; whether the relationship generalizes to other genomes, including those with substantially different TF-family compositions, is untested. Finally, our literature search, while conducted specifically and iteratively against this result (Introduction, Discussion), cannot rule out obscure or non-indexed prior work, and a dedicated, exhaustive review remains advisable before any formal submission.

## 5. Methods

### Data sources

STRING v12.0 D. melanogaster protein annotation, combined-score network, and full evidence-channel-split network (taxon 7227) [20,21,22]; JASPAR CORE Drosophila position frequency matrices [23]; JASPAR2026 dm6 genome-wide predicted binding sites (bigBed); 124-way phyloP conservation scores for dm6 [24,25,26]; the Ensembl BDGP6.54 / release 6 D. melanogaster gene annotation (GTF) [27]; and the FlyBase gene expression RPKM matrix (167 conditions) [28,29].

### TF identification and family classification

Candidate TFs were flagged from STRING protein annotations by a DNA-binding-domain keyword regular expression (zinc finger, homeobox, bHLH, bZIP, forkhead, nuclear receptor, and related terms, deliberately including structural keywords beyond the bare phrase “transcription factor” to reduce annotation-text circularity), then confirmed by name-matching against JASPAR Drosophila motifs, yielding 279–304 confirmed TFs depending on downstream filtering. DNA-binding-domain family was assigned independently via a second annotation-keyword regular expression spanning 12 major families plus an “other/unclassified” and “multiple” category.

### PPI network construction and evidence-channel decomposition

The combined-score network was deduplicated to undirected edges (i < j). The physical-evidence network was built from STRING’s full evidence-channel file by combining the experiments, experiments_transferred, database, and database_transferred columns via STRING’s standard probabilistic-OR formula, 1 − ∏(1 − p_i_), explicitly excluding neighborhood, fusion, cooccurrence, homology, coexpression, and textmining channels. Module fraction for a given TF was defined as the fraction of its network degree (in the relevant network) directed at other confirmed TFs.

### Genome-wide binding-site density

Two independent ∼18 Mb genomic samples were constructed from the GTF: a gene-body-restricted sample (protein-coding gene spans, capped at 40 kb per gene to prevent domination by outlier loci) and an intergenic-restricted sample (gaps between genes, with a 2 kb buffer excluded on each side, similarly capped). JASPAR2026 bigBed entries and phyloP values were fetched over both samples using pyBigWig, with checkpointed, resumable aggregation into per-TF scalar accumulators. Following the diagnosis described in Results, no PWM match-score threshold was applied when counting predicted sites, since any fixed absolute threshold discards a motif-length-dependent fraction of sites and reintroduces a cross-motif comparability artifact.

### Expression breadth and composite index

Expression breadth was computed from the FlyBase RPKM matrix as the fraction of conditions with RPKM > 1 per TF. The composite specialization index averaged rank-z-transformed module fraction and rank-z-transformed inverse expression breadth.

### Statistical framework

All partial correlations used rank-z-transformed variables with linear OLS residualization on the named control set, followed by Pearson correlation of the residuals. TF–TF assortativity used both naive random-relabeling permutation nulls and degree-matched permutation nulls (relabeling within degree bins, 5–50 bins tested), 500–1,000 permutations each. The family-cluster bootstrap resampled whole DNA-binding-domain families with replacement, 1,500–2,000 iterations. Multiple-testing correction used the Benjamini-Hochberg procedure [35] across the core battery of tests. All analysis code is provided in full (Data and Code Availability) and was independently re-executed end-to-end against the raw input files as part of this study’s verification process, including a full, successful dry run that reproduced every reported figure directly from source data.

**Table 1.** Overview of the fourteen-stage analysis pipeline.

| Stage | Purpose |
| --- | --- |
| 1–2 | Identify TFs from STRING annotation (DNA-binding-domain keyword regex) and parse JASPAR PFMs into per-position Shannon information content. |
| 3–4 | Match JASPAR motifs to STRING protein IDs (304 confirmed TFs); build the combined-score PPI network; classify DNA-binding-domain family by an independent annotation-keyword regex. |
| 5–6 | Compute expression breadth and pairwise co-expression from FlyBase RPKM; construct two independent ~18 Mb genomic samples (gene-body-restricted and intergenic-restricted) from the Ensembl GTF. |
| 7–8 | Fetch JASPAR2026 genome-wide predicted sites and phyloP conservation over both samples (checkpointed, unfiltered — no PWM score threshold, to avoid the motif-length artifact described in Results); compare genic vs. intergenic compartments. |
| 9–10 | Decompose STRING evidence channels; retain only experimental and curated-database evidence (excluding textmining, co-expression, genomic-context, and homology channels) to build a physical-evidence-only PPI network; compute the core module-fraction vs. site-density test. |
| 11 | Composite specialization index (module fraction + inverse expression breadth); TF–TF assortativity via naive and degree-matched permutation nulls (500–1,000 permutations). |
| 12–13 | Confound battery (motif length, GC/AT composition, family/paralog exclusion) and TF-family cluster bootstrap (2,000 resamples). |
| 14 | Adversarial robustness battery: tautology/collinearity check, Spearman vs. Pearson, outlier leverage trimming, winsorizing, post-hoc power. |

## 6. Data and Code Availability

The complete analysis pipeline ([github-hyperlink](https://github.com/ramanujan2710/Interactome-Specialization-Predicts-Genome-Wide-Binding-Site-Degeneracy-)) is provided as a standalone, staged, checkpointed Python script that reproduces every result and figure in this manuscript end-to-end from the raw STRING, JASPAR, phyloP, GTF, and FlyBase input files listed above. The script is organized into 14 sequential, independently re-runnable stages corresponding to the Methods subsections above, and includes inline documentation of the score-threshold pitfall described in Results. Figures were generated directly from pipeline output tables using matplotlib and are fully reproducible from the same intermediate files the pipeline writes to disk.

## Notes

https://github.com/ramanujan2710/Interactome-Specialization-Predicts-Genome-Wide-Binding-Site-Degeneracy-

### Competing Interest Statement

The authors have declared no competing interest.

## References

1. Schneider TD, Stormo GD, Gold L, Ehrenfeucht A. Information content of binding sites on nucleotide sequences. J Mol Biol. 1986;188(3):415–431.

2. Schneider TD, Stephens RM. Sequence logos: a new way to display consensus sequences. Nucleic Acids Res. 1990;18(20):6097–6100.

3. Stormo GD. Modeling the specificity of protein-DNA interactions. Quant Biol. 2013;1(2):115–130.

4. Zhang C, Xuan Z, Otto S, Hover JR, McCorkle SR, Mandel G, Zhang MQ. A clustering property of highly-degenerate transcription factor binding sites in the mammalian genome. Nucleic Acids Res. 2006;34(8):2238–2246.

5. Savir Y, Kagan J, Tlusty T. Binding of transcription factors adapts to resolve information-energy trade-off. arXiv:1505.01215 [q-bio.MN]. 2015.

6. Mahmutovic A, Berg OG, Elf J. Speed-specificity trade-offs in the transcription factors search for their genomic binding sites. Trends Genet. 2021;37(5):421–432.

7. Wunderlich Zn, Mirny LA. Different gene regulation strategies revealed by analysis of binding motifs. Trends Genet. 2009;25(10):434–440.

8. Cirillo D, Botta-Orfila T, Tartaglia GG. By the company they keep: interaction networks define the binding ability of transcription factors. Nucleic Acids Res. 2015;43(19):e125.

9. Rhee DY, Cho DY, Zhai B, Slattery M, Ma L, Mintseris J, Wong CY, White KP, Celniker SE, Przytycka TM, Gygi SP, Obar RA, Artavanis-Tsakonas S. Transcription factor networks in Drosophila melanogaster. Cell Rep. 2014;8(6):2031–2043.

10. Shokri L, Inukai S, Hafner A, et al. A comprehensive Drosophila melanogaster transcription factor interactome. Cell Rep. 2019;27(3):955–970.e7.

11. Naderi J, Magalhaes AP, Kibar G, Stik G, Zhang Y, Mackowiak SD, Wieler HM, Rossi F, Buschow R,Christou-Kent M, Alcoverro-Bertran M, Graf T, Vingron M, Hnisz D. An activity-specificity trade-off encoded in human transcription factors. Nat Cell Biol. 2024;26:1309–1321.

12. Ji W, Hachmo O, Barkai N, Amir A. Design principles of transcription factors with intrinsically disordered regions. eLife. 2025;13:RP104956 (preprint: bioRxiv 2024.11.26.625383).

13. Boija A, Klein IA, Sabari BR, Dall’Agnese A, Coffey EL, Zamudio AV, Li CH, Shrinivas K, Manteiga JC, Hannett NM, Abraham BJ, Afeyan LK, Guo YE, Rimel JK, Fant CB, Schuijers J, Lee TI, Taatjes DJ, Young RA. Transcription factors activate genes through the phase-separation capacity of their activation domains. Cell. 2018;175(7):1842–1855.

14. Sabari BR, Dall’Agnese A, Boija A, Klein IA, Coffey EL, Shrinivas K, Abraham BJ, Hannett NM, Zamudio AV, Manteiga JC, Li CH, Guo YE, Day DS, Schuijers J, Vasile E, Malik S, Hnisz D, Lee TI, Cisse II, Roeder RG, Sharp PA, Chakraborty AK, Young RA. Coactivator condensation at super-enhancers links phase separation and gene control. Science. 2018;361(6400):eaar3958.

15. Lukatsky DB, Shakhnovich EI. Sequence correlations shape protein promiscuity. arXiv:1004.5048 [q-bio.BM]. 2010.

16. Melamed D, Young DL, Miller CR, Fields S. Protein interaction evolution from promiscuity to specificity with reduced flexibility in an increasingly complex network. Sci Rep. 2017;7:44948.

17. Jeong H, Mason SP, Barabasi AL, Oltvai ZN. Lethality and centrality in protein networks. Nature. 2001;411:41–42.

18. Newman MEJ. Assortative mixing in networks. Phys Rev Lett. 2002;89(20):208701.

19. Maslov S, Sneppen K. Specificity and stability in topology of protein networks. Science. 2002;296(5569):910–913.

20. von Mering C, Huynen M, Jaeggi D, Schmidt S, Bork P, Snel B. STRING: a database of predicted functional associations between proteins. Nucleic Acids Res. 2003;31(1):258–261.

21. Szklarczyk D, Kirsch R, Koutrouli M, et al. The STRING database in 2023: protein-protein association networks and functional enrichment analyses for any sequenced genome of interest. Nucleic Acids Res. 2023;51(D1):D638–D646.

22. Szklarczyk D, Nastou K, Koutrouli M, et al. The STRING database in 2025: protein networks with directionality of regulation. Nucleic Acids Res. 2025;53(D1):D730–D737.

23. Rauluseviciute I, Riudavets-Puig R, Blanc-Mathieu R, Castro-Mondragon JA, Ferenc K, Kumar V, Lemma RB, Lucas J, Cheneby J, Baranasic D, Khan A, Fornes O, Gundersen S, Johansen M, Hovig E, Lenhard B, Sandelin A, Wasserman WW, Parcy F, Mathelier A. JASPAR 2024: 20th anniversary of the open-access database of transcription factor binding profiles. Nucleic Acids Res. 2024;52(D1):D174–D182.

24. Siepel A, Bejerano G, Pedersen JS, Hinrichs AS, Hou M, Rosenbloom K, Clawson H, Spieth J, Hillier LW, Richards S, Weinstock GM, Wilson RK, Gibbs RA, Kent WJ, Miller W, Haussler D. Evolutionarily conserved elements in vertebrate, insect, worm, and yeast genomes. Genome Res. 2005;15(8):1034–1050.

25. Pollard KS, Hubisz MJ, Rosenbloom KR, Siepel A. Detection of nonneutral substitution rates on mammalian phylogenies. Genome Res. 2010;20(1):110–121.

26. Hubisz MJ, Pollard KS, Siepel A. PHAST and RPHAST: phylogenetic analysis with space/time models. Brief Bioinform. 2011;12(1):41–51.

27. dos Santos G, Schroeder AJ, Goodman JL, et al. FlyBase: introduction of the Drosophila melanogaster Release 6 reference genome assembly and large-scale migration of genome annotations. Nucleic Acids Res. 2015;43(D1):D690–D697.

28. Gramates LS, Agapite J, Attrill H, et al. (FlyBase Consortium). FlyBase: updates to the Drosophila genes and genomes database. Genetics. 2024;227(1):iyad211.

29. Larkin A, Marygold SJ, Antonazzo G, et al. (FlyBase Consortium). FlyBase: updates to the Drosophila melanogaster knowledge base. Nucleic Acids Res. 2021;49(D1):D899–D907.

30. Spitz F, Furlong EEM. Transcription factors: from enhancer binding to developmental control. Nat Rev Genet. 2012;13(9):613–626.

31. Zinzen RP, Girardot C, Gagneur J, Braun M, Furlong EEM. Combinatorial binding predicts spatio-temporal cis-regulatory activity. Nature. 2009;462(7269):65–70.

32. Zaret KS, Carroll JS. Pioneer transcription factors: establishing competence for gene expression. Genes Dev. 2011;25(21):2227–2241.

33. Li XY, Thomas S, Sabo PJ, Eisen MB, Stamatoyannopoulos JA, Biggin MD. The role of chromatin accessibility in directing the widespread, overlapping patterns of Drosophila transcription factor binding. Genome Biol. 2011;12(4):R34.

34. Roy S, Ernst J, Kharchenko PV, et al. (modENCODE Consortium). Identification of functional elements and regulatory circuits by Drosophila modENCODE. Science. 2010;330(6012):1787–1797.

35. Benjamini Y, Hochberg Y. Controlling the false discovery rate: a practical and powerful approach to multiple testing. J R Stat Soc Series B. 1995;57(1):289–300.

36. Duret L, Mouchiroud D. Determinants of substitution rates in mammalian genes: expression pattern affects selection intensity but not mutation rate. Mol Biol Evol. 2000;17(1):68–74.

37. Winter EE, Goodstadt L, Ponting CP. Elevated rates of protein secretion, evolution, and disease among tissue-specific genes. Genome Res. 2004;14(1):54–61.

38. Johnson AF, Nguyen HT, Veitia RA. Causes and effects of haploinsufficiency. Biol Rev Camb Philos Soc. 2019;94(5):1774–1785.

39. Naqvi S, Kim S, Hoskens H, Matthews HS, Spritz RA, Klein OD, Hallgrimsson B, Swigut T, Claes P, Pritchard JK, Wysocka J. Precise modulation of transcription factor levels identifies features underlying dosage sensitivity. Nat Genet. 2023;55:841–851.

40. Bonchuk A, Boyko K, Fedotova A, Nikolaeva A, Lushchekina S, Khrustaleva A, Popov V, Georgiev P. Structural basis of diversity and homodimerization specificity of zinc-finger-associated domains in Drosophila. Nucleic Acids Res. 2021;49(4):2375–2389.

41. Vinayagam A, Stelzl U, Foulle R, Plassmann S, Zenkner M, Timm J, Assmus HE, Andrade-Navarro MA, Wanker EE. A directed protein interaction network for investigating intracellular signal transduction. Sci Signal. 2011;4(189):rs8.

42. Ravasi T, Suzuki H, Cannistraci CV, et al. An atlas of combinatorial transcriptional regulation in mouse and man. Cell. 2010;140(5):744–752.

43. Reece-Hoyes JS, Diallo A, Lajoie B, Kent A, Shrestha S, Kadreppa S, Pesyna C, Dekker J, Myers CL, Walhout AJM. Enhanced yeast one-hybrid assays for high-throughput gene-centered regulatory network mapping. Nat Methods. 2011;8(12):1059–1064.

44. Kribelbauer JF, Rastogi C, Bussemaker HJ, Mann RS. Low-affinity binding sites and the transcription factor specificity landscape. Annu Rev Cell Dev Biol. 2019;35:357–379.

45. Lambert SA, Jolma A, Campitelli LF, Das PK, Yin Y, Albu M, Chen X, Taipale J, Hughes TR, Weirauch MT. The human transcription factors. Cell. 2018;172(4):650–665.

46. Slattery M, Riley T, Liu P, Abe N, Gomez-Alcala P, Dror I, Zhou T, Rohs R, Honig B, Bussemaker HJ, Mann RS. Cofactor binding evokes latent differences in DNA binding specificity between Hox proteins. Cell. 2011;147(6):1270–1282.

